# HSD17B11 maintains FSP1 localization on lipid droplets to support ferroptosis defense

**DOI:** 10.64898/2026.05.19.726342

**Authors:** Valeria Montenegro Vazquez, Baley A. Goodson, Oralia M. Kolaczkowski, Halima Akter, Li Chen, Nikesh Narang, Jaya Rajaiya, Monica Rosas Lemus, Jing Pu

**Affiliations:** Department of Molecular Genetics and Microbiology, University of New Mexico Health Sciences Center, Albuquerque, NM, USA; Autophagy, Inflammation, & Metabolism Center of CoBRE, University of New Mexico Health Sciences Center, Albuquerque, NM, USA; Department of Ophthalmology & Visual Sciences, University of New Mexico Health Sciences Center, Albuquerque, NM, USA

**Keywords:** Ferroptosis, Lipid metabolism, N-myristoylation, Organelle biology, Cell death

## Abstract

Ferroptosis is a regulated form of cell death driven by iron-dependent lipid peroxidation, and lipid droplets (LDs) are increasingly recognized as important regulators of this process. Consistent with this role, the anti-ferroptotic factor ferroptosis suppressor protein 1 (FSP1) is known localizing on LDs through N-myristoylation-dependent membrane targeting, where it protects LD lipids from peroxidation. Here, we identify the LD protein HSD17B11 as an additional factor required for maintaining both FSP1 localization on LDs and cellular FSP1 abundance. Silver staining followed by mass spectrometry analysis of purified LD proteins identified reduced LD-associated FSP1 in HSD17B11-deficient cells, which was further validated by immunoblotting and imaging analyses. Mechanistically, HSD17B11 physically interacted with FSP1 and was required to preserve FSP1 association with LDs. Mutational analyses further demonstrated that both FSP1 N-myristoylation and an intact HSD17B11 interaction interface are necessary for LD targeting. Correspondingly, HSD17B11 deficiency reduced LD-associated and total cellular FSP1 levels and increased cellular sensitivity to lipid oxidative stress. Together, our findings identify HSD17B11 as a previously unrecognized regulator of LD-associated FSP1 and reveal an additional mechanism controlling compartmentalized ferroptosis defense.

## INTRODUCTION

Ferroptosis is a regulated form of cell death, driven by iron-dependent lipid peroxidation^1^, and has emerged as an important mechanism contributing to cancer progression^2,3^, acute tissue injury^4^, and neurodegenerative diseases^5^. Unlike apoptosis or necrosis, ferroptosis is characterized by the accumulation of oxidized polyunsaturated phospholipids, which ultimately compromise membrane integrity and cellular viability^6,7^. Because uncontrolled lipid peroxidation is highly toxic, cells have evolved multiple defense systems to suppress ferroptosis and maintain lipid redox homeostasis. Understanding these protective mechanisms may reveal therapeutic targets for diseases associated with oxidative stress and ferroptotic injury.

Among the known ferroptosis defense pathways, glutathione peroxidase 4 (GPX4)^8^ and ferroptosis suppressor protein 1 (FSP1, also known as AIFM2)^9,10^ function as two major anti-ferroptotic systems. GPX4 reduces phospholipid hydroperoxides using glutathione as a cofactor and is considered a central regulator of ferroptosis resistance^8^. In parallel, FSP1 suppresses ferroptosis independently of GPX4 by reducing coenzyme Q10 to generate lipophilic radical-trapping antioxidants^9,10^.

Although GPX4 and FSP1 suppress lipid peroxidation at cellular membranes, increasing evidence suggests that intracellular organelles also actively participate in regulating ferroptosis sensitivity. Lipid droplets (LDs), central organelle in lipid metabolism, are increasingly recognized as important modulators of ferroptosis^6,7^. LDs can protect cells by sequestering PUFAs into neutral lipids, thereby limiting their incorporation into peroxidation-sensitive membranes^11,12^. In addition, LDs dynamically interact with mitochondria^13,14^ and peroxisomes^15,16^ to support fatty acid oxidation and lipid remodeling, suggesting that LD-associated metabolism is an important determinant of ferroptotic sensitivity.

Unlike the broadly distributed GPX4 system, FSP1 localizes to both the plasma membrane and LDs^9,10^, further suggesting that LDs function as intracellular hubs for controlling lipid peroxidation. Consistent with this model, recent lipidomics analyses showed that loss of FSP1 promotes neutral lipid peroxidation within LDs and initiates ferroptosis^17^. These observations suggest that the protective function of FSP1 depends not only on its enzymatic activity, but also on its proper membrane localization.

FSP1 is targeted to membranes through N-myristoylation at its N-terminus^9,10^, and pharmacological displacement of FSP1 from membranes induces cytosolic phase separation and cell death^18^, highlighting the importance of membrane association in regulating FSP1 stability and anti-ferroptotic activity.

Hydroxysteroid 17-beta dehydrogenases (HSD17Bs) are members of the short-chain dehydrogenase/reductase superfamily and participate in diverse aspects of steroid and lipid metabolism^19^. Several HSD17B family proteins associate with LDs with broader functions in organizing LD-associated metabolic processes. Among them, HSD17B13 is a liver-enriched LD protein genetically linked to protection against metabolic liver disease^20^. HSD17B11 is a closely related paralog of HSD17B13 that also localizes to LDs and broadly expresses across multiple tissues and organs^19^, but its physiological functions remain poorly understood.

Here, we identify HSD17B11 as a critical regulator of LD-associated FSP1. Loss of HSD17B11 disrupts FSP1 localization to LDs and sensitizes cells to lipid oxidative stress. Mechanistically, HSD17B11 physically interacts with FSP1 and is required for maintaining FSP1 association with LDs. Our findings uncover a previously unrecognized mechanism linking LD biology to ferroptosis defense and establish HSD17B11 as a key regulator of compartmentalized antioxidant protection.

## RESULTS

### HSD17B11 deficiency changed lipid droplet morphology and protein composition

To investigate the role of HSD17B11 in lipid droplet (LD) biology, we generated a knockout (KO) in the human hepatocellular carcinoma cell line Huh7 using CRISPR-Cas9. Immunofluorescence confirmed the successful deletion, showing loss of HSD17B11 from LD surfaces in KO cells, while the LD marker perilipin 2 (PLIN2, also known as ADRP) remained intact (Fig. 1A), indicating preserved LD structure. Immunoblotting further confirmed the depletion of HSD17B11 without significant changes in other LD-associated proteins, including its paralog HSD17B13 and PLIN family members PLIN2 and PLIN3 (also known as TIP47) (Fig. 1B).

**Figure 1.**
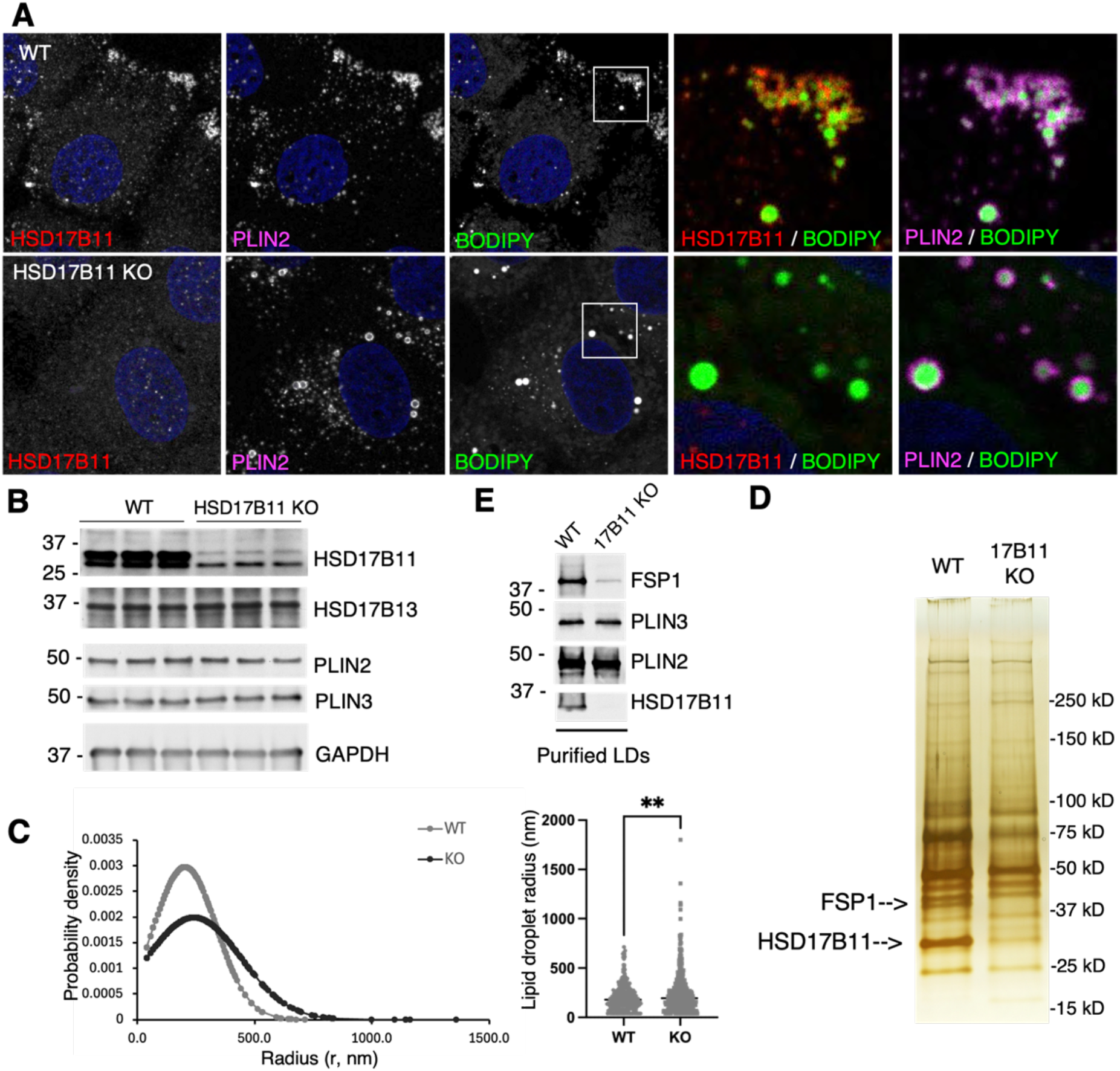
Characterization of lipid droplets in HSD17B11-knockout (KO) cells. **A**. Immunofluorescence and confocal microscopy analysis of wild-type (WT) and HSD17B11-KO Huh7 cells. **B**. Immunoblotting analysis of whole-cell lysates from WT and HSD17B11-KO cells. **C**. Individual lipid droplet radii were measured using ImageJ and plotted as probability density distributions using Microsoft Excel. Data were pooled from four cells per group. Statistical significance was determined using Welch’s *t* test. **, *p*=0.0014. **D**. Proteins extracted from purified lipid droplets were analyzed by silver staining. Bands indicated by arrows were excised and subjected to mass spectrometry for protein identification. **E**. Immunoblotting analysis of purified LD fractions from WT and HSD17B11-KO cells.

Interestingly, HSD17B11 deletion altered LD morphology and distribution. KO cells appeared to contain fewer peripherally distributed LDs (Fig. 1A) and a significant increase in LD size (Fig. 1C), suggesting that HSD17B11 plays a fundamental role in regulating LD organization.

To determine whether HSD17B11 affects LD protein composition, we purified LDs from WT and KO cells. Silver staining revealed two protein bands present in WT but absent in KO LD fractions. Mass spectrometry identified these bands as HSD17B11 and ferroptosis suppressor protein 1 (FSP1) (Fig. 1D). To validate this result, we performed immunoblotting on purified LD fractions. While PLIN2 and PLIN3 levels remained largely unchanged, FSP1 was markedly reduced in KO LDs (Fig. 1D). These findings suggest a previously unrecognized role for HSD17B11 in regulating ferroptosis through modulation of LD-associated FSP1.

### FSP1 dissociates from lipid droplets upon HSD17B11 loss

Immunofluorescence analysis confirmed the loss of LD FSP1 in HSD17B11-deficient cells (Fig. 2A), which was further supported by high-content imaging quantification (Fig. 2B). Similarly, siRNA-mediated knockdown of HSD17B11 recapitulated the reduction of LD-associated FSP1 (Fig. 2C). Importantly, re-expression of HSD17B11 in KO cells restored LD FSP1, whereas expression of HSD17B13 failed to do so (Fig. 2D). These results demonstrate that HSD17B11 is specifically required for maintaining LD FSP1, independent of potential CRISPR off-target effects.

**Figure 2.**
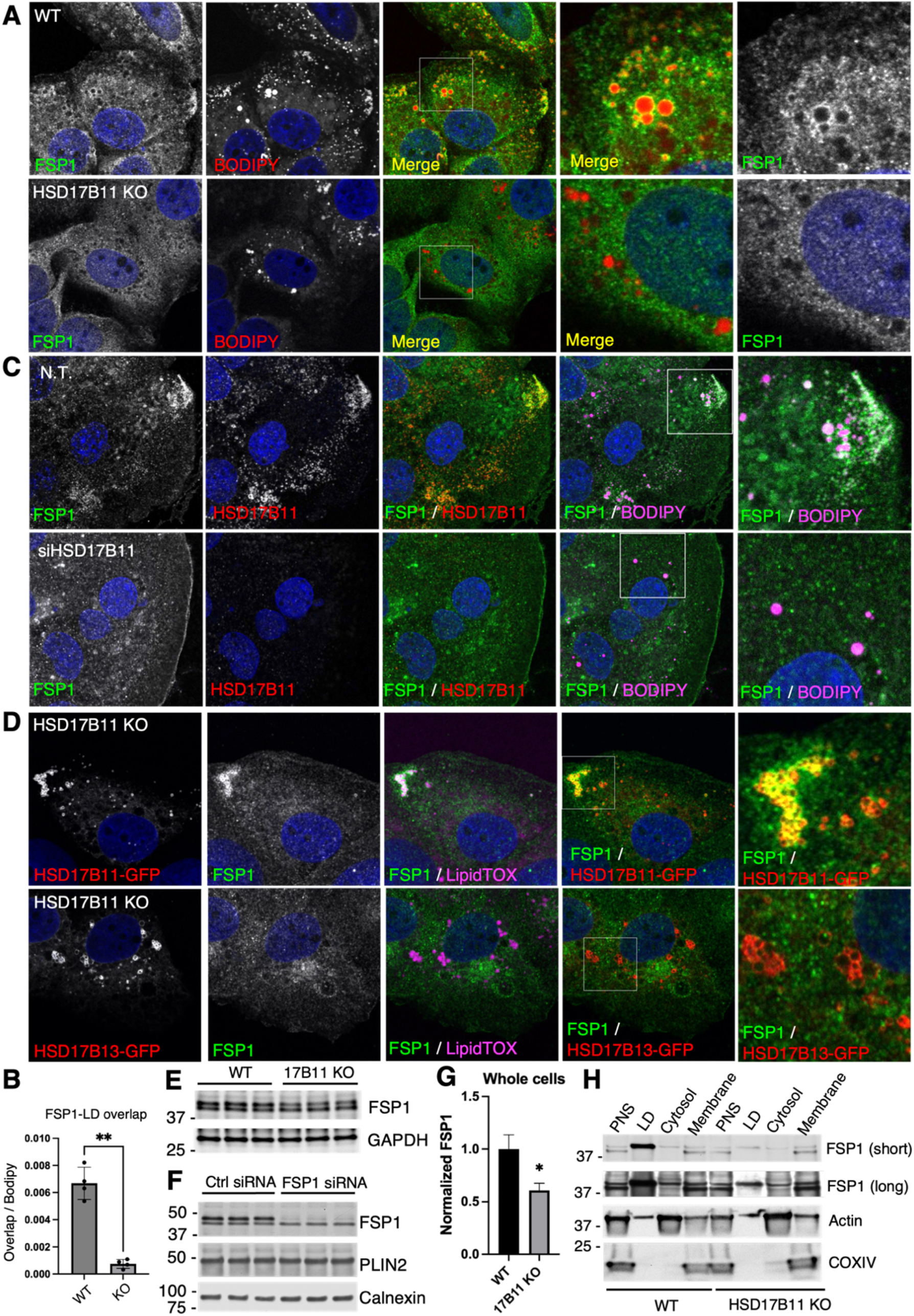
Characterization of FSP1 localization and expression. **A**,**B**. Immunofluorescence analysis of wildtype (WT) and HSD17B11-knockout (KO) Huh7 cells imaged by confocal microscopy (A) and quantified using a high-content imaging system (B). **C**. Immunofluorescence and confocal microscopy analysis of cells transfected with non-targeting (N.T.) or HSD17B11 siRNA. **D**. HSD17B11-KO cells were transfected with HSD17B11 or HSD17B13 constructs followed by immunofluorescence and confocal microscopy analysis. **E**,**F**. Immunoblotting analysis of whole-cell lysates from the indicated cells. **G**. Quantification of FSP1 bands shown in (F) using ImageJ. Data are presented as mean ± SD. Statistical significance was determined using Student’s *t* test. *, *p* < 0.05. **H**. Post-nuclear supernatants (PNS) were fractionated into lipid droplet (LD), cytosolic, and total membrane fractions followed by immunoblotting with the indicated antibodies.

We next asked whether the reduction of LD-associated FSP1 was due to decreased total protein levels or redistribution. Immunoblotting of whole-cell lysates revealed two bands near the expected molecular weight of FSP1 (Fig. 2E). siRNA-mediated knockdown selectively reduced the upper band, confirming it as FSP1, whereas the lower band remained unchanged and is therefore likely non-specific (Fig. 2F). Quantification of the FSP1 bands showed an approximately 40% reduction in total FSP1 levels in HSD17B11-deficient cells (Fig. 2G), which is less pronounced than the depletion observed in LD fractions (Fig. 1E), suggesting that loss of LD-associated FSP1 cannot be explained solely by reduced expression.

To determine whether FSP1 is redistributed, we fractionated cells into LD, cytosolic, and membrane compartments. In HSD17B11-KO cells, FSP1 was strongly reduced in the LD fraction and modestly reduced in the cytosol (Fig. 2H), likely contributing to the decreased total cellular FSP1 levels (Fig. 2E,G). In contrast, FSP1 levels in membrane fractions remained unchanged compared to WT cells (Fig. 2H), suggesting that plasma membrane-associated FSP1 was preserved. Notably, the loss of FSP1 from LDs was not accompanied by a corresponding increase in other cellular fractions (Fig. 2H), suggesting that dissociated FSP1 is destabilized and subsequently degraded. Consistent with this interpretation, recent studies demonstrated that FSP1 is subject to ubiquitin-mediated proteasomal degradation^21^. Together, these results indicate that HSD17B11 loss leads to dissociation of FSP1 from LDs, while plasma membrane-associated FSP1 remains largely unaffected.

### FSP1 physically interacts with HSD17B11

HSD17B11 associates with LD membranes via its N-terminal helix^22^, while FSP1 is targeted to membranes through N-myristoylation^9,10^. Structural modeling using AlphaFold predicted that the N-terminal regions of both proteins orient in the same direction at the membrane interface (Fig. 3A), consistent with their membrane-binding mechanisms.

**Figure 3.**
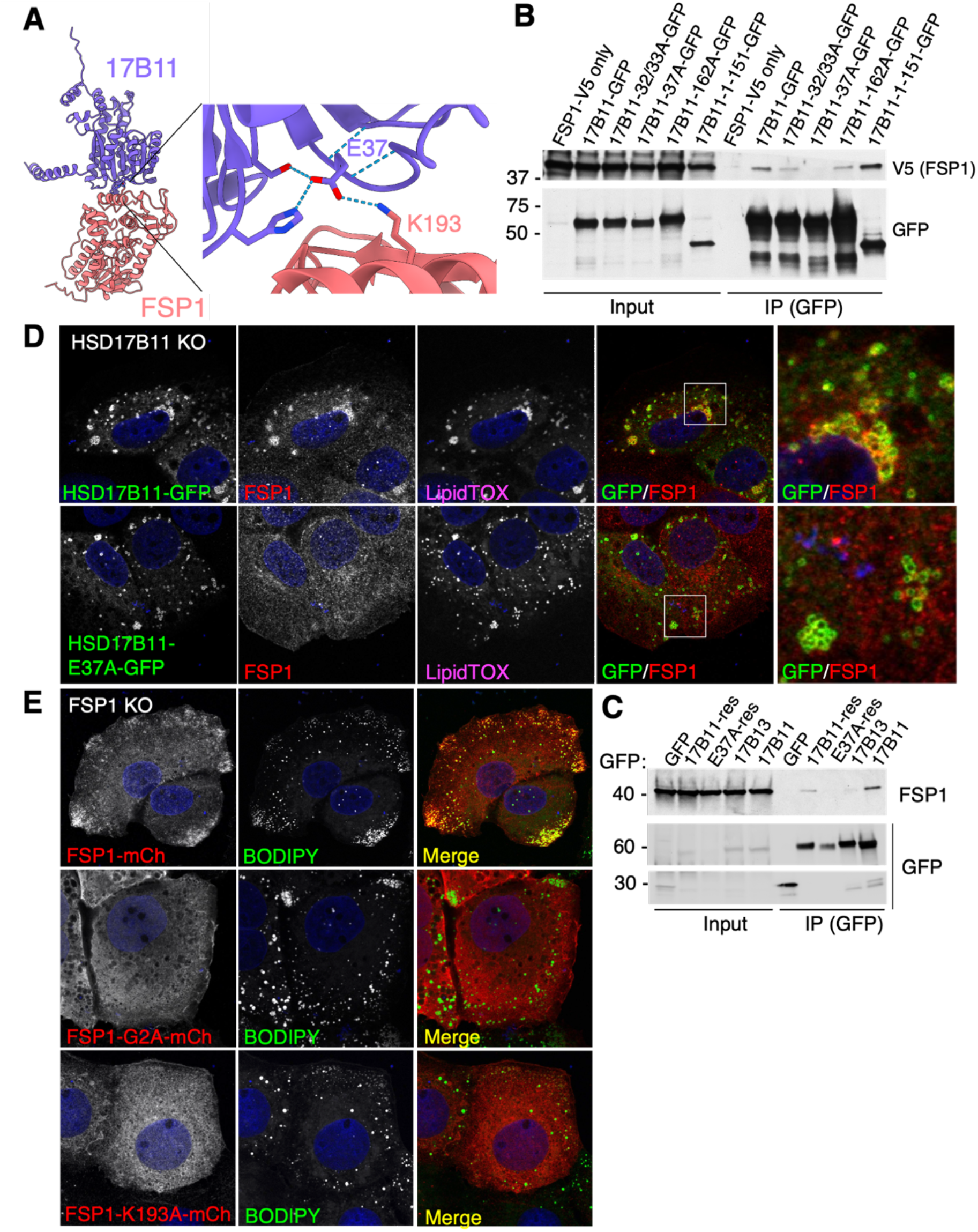
Identification of HSD17B11-FSP1 interaction interface. **A**. AlphaFold-predicted interaction model of human HSD17B11 and FSP1. Dotted lines indicate predicted hydrogen bonds. **B**. Co-immunoprecipitation (co-IP) analysis of transfected HSD17B11-GFP WT, indicated mutants, or the shorter HSD17B11 isoform with FSP1-V5. **C**. GFP-fused WT HSD17B11 or the E37A mutant was stably expressed in HSD17B11-KO cells, followed by co-IP analysis to detect endogenous FSP1. Co-IP analysis of transiently transfected HSD17B11 and HSD17B13 was also performed for comparison. **D.E**, Immunofluorescence and confocal microscopy analysis of cells transfected with the indicated constructs.

To test this interaction model, we generated point mutations in HSD17B11 at predicted interface residues and performed co-immunoprecipitation (co-IP) assays. Wild-type HSD17B11 efficiently co-immunoprecipitated FSP1, whereas mutation of glutamic acid 37 to alanine (E37A) abolished this interaction (Fig. 3B). Other mutations partially reduced binding (Fig. 3B), suggesting that multiple residues contribute to the interaction interface. A truncated HSD17B11 isoform containing residues 1–151 retained and even enhanced binding to FSP1 (Fig. 3B), indicating that the N-terminus is both necessary and sufficient for interaction. Consistently, stable expression of GFP-tagged WT HSD17B11 in KO cells restored interaction with endogenous FSP1, whereas the E37A mutant did not (Fig. 3C). In contrast to HSD17B11, transiently expressed HSD17B13 also failed to interact with FSP1 (Fig. 3C), consistent with its inability to rescue FSP1 localization to LDs (Fig. 2D).

Reciprocally, WT or mutant FSP1 constructs were expressed in FSP1-KO cells to assess LD localization. WT FSP1 localized robustly to LDs, whereas the K193A mutant, which is predicted to disrupt interaction with HSD17B11, failed to localize to LDs (Fig. 3E). Similarly, the G2A mutant, which abolishes N-myristoylation, also lost LD association (Fig. 3E). These results support that FSP1 localization to LDs requires both N-myristoylation and an intact interface with HSD17B11.

### HSD17B11 stabilizes FSP1 on LDs and protects cells from ferroptosis

We further tested whether HSD17B11 deficiency affected FSP1 dissociation from LDs through disruption of the HSD17B11–FSP1 interaction. In the cell fractionation assay, expression of WT HSD17B11 restored LD-associated FSP1 in KO cells, whereas the E37A mutant failed to do so (Fig. 4A), indicating that the interaction is required for FSP1 localization. These results support a model in which HSD17B11 recruits and stabilizes FSP1 on LDs through direct protein-protein interaction.

**Figure 4.**
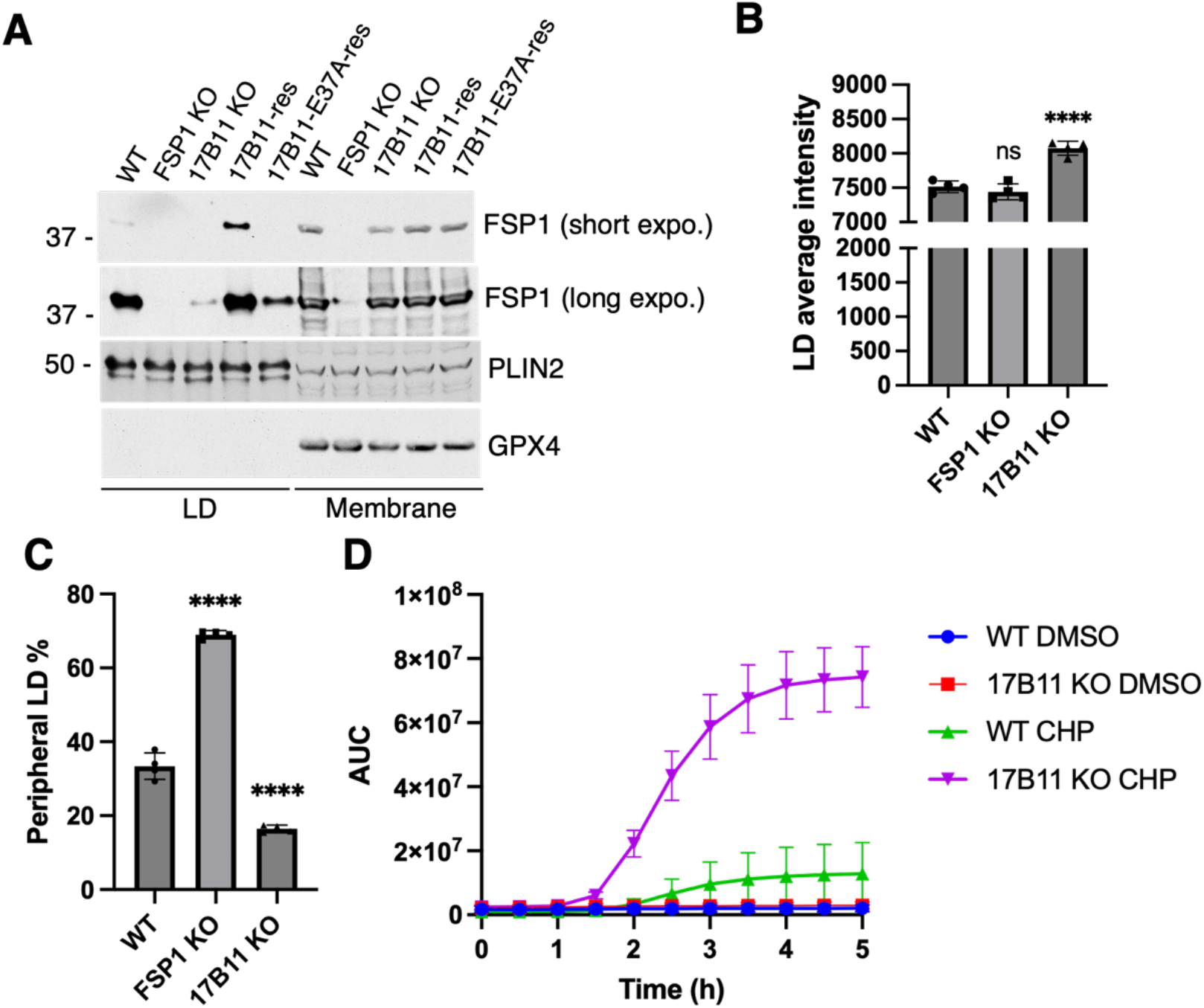
HSD17B11 stabilizes LD-associated FSP1 and protects cells from ferroptotic stress. **A**. Lipid droplet (LD) and total membrane fractions from the indicated cells were analyzed by immunoblotting with the indicated antibodies. **B.C**, LD morphology and distribution were analyzed using a high-content imaging system. More than 1,000 cells per well and four wells per cell line were analyzed. Data are presented as mean ± SD at the well level. Statistical significance was determined using one-way ANOVA. ****, *p* < 0.0001; ns, not significant. **D**. Cell death analysis of the indicated cells treated with DMSO or 50 μM cumene hydroperoxide (CHP) to induce ferroptotic stress. Fluorescence was monitored using the Incucyte live-cell imaging system.

To test whether HSD17B11 deficiency-caused LD morphology and positioning alteration is related to loss of FSP1 on LDs, we compared LD features among the WT, HSD17B11-KO, and FSP1-KO cells. Contrast to HSD17B11-KO cells, FSP1-KO cells did not significantly alter LD size or reduce cell periphery LD distribution (Fig. 4B,C), indicating that the LD feature alteration was not attributed to FSP1 LD localization.

Considering the known anti-ferroptotic function of FSP1, we tested cell ferroptosis sensitivity in WT and HSD17B11-KO cells. Induction of ferroptosis by using a lipid-targeting oxidative stress trigger, cumene hydroperoxide (CHP), markedly enhanced cell death in the HSD17B11-KO cells (Fig. 4D). Since HSD17B11 deletion only affected LD but plasma membrane FSP1, these results highlight the role of LDs in protecting cells from oxidation stress. Together, this work defines HSD17B11 as a multifunctional protein on LDs, and its anti-ferroptosis function is through localizing FSP1 on LDs.

## DISCUSSION

Our study identifies HSD17B11 as a previously unrecognized regulator of lipid droplet (LD)-associated ferroptosis defense through maintaining FSP1 localization on LDs. Although FSP1 has emerged as a major ferroptosis suppressor and an attractive therapeutic target in cancer^18,23^, the mechanisms regulating its intracellular localization and stability remain incompletely understood. Previous studies primarily focused on N-myristoylation-mediated membrane targeting as the key determinant of FSP1 function^9,10^. More recently, vitamin B2 metabolism was shown to support FSP1 stability^21^. Here, we demonstrate that LD localization and cellular abundance of FSP1 additionally require HSD17B11, revealing a distinct layer of regulation.

Mechanistically, HSD17B11 physically interacts with FSP1 and is required for stable FSP1 association with LDs. Loss of HSD17B11 resulted in dissociation of FSP1 from LDs and reduction of FSP1 protein levels, suggesting that LD association contributes to FSP1 stability. These findings expand the current understanding of FSP1 regulation and identify HSD17B11 as a potential target for manipulating ferroptosis sensitivity.

FSP1 has attracted substantial interest as a therapeutic target because ferroptosis resistance contributes to tumor survival and therapy resistance. Multiple small-molecule FSP1 inhibitors have been developed to sensitize cancer cells to ferroptosis^18,24,25^. Our findings suggest an alternative strategy for modulating FSP1 function through disrupting its LD localization and stability. Because LD localization is critical for the anti-ferroptotic function of FSP1^17^, mechanisms controlling its compartmentalization may represent an additional level of ferroptosis regulation. Future studies will be needed to determine whether HSD17B11-mediated FSP1 localization and stability can be therapeutically targeted and whether this pathway contributes to ferroptosis resistance *in vivo*.

Our results also strengthen the emerging concept that LDs function as active regulators of lipid peroxidation and ferroptosis rather than passive lipid storage organelles. As the central organelles for neutral lipid storage and fatty acid buffering, LDs are highly sensitive to changes in lipid flux and metabolic homeostasis. Increasing evidence suggests that LDs can protect cells by sequestering polyunsaturated fatty acids away from peroxidation-sensitive membrane phospholipids^11,26^. Consistent with this idea, recent lipidomics studies demonstrated that loss of FSP1 induces neutral lipid peroxidation within LDs before widespread ferroptotic damage occurs^17^. Together with these observations, our findings support a model in which LDs act as early sensors and regulators of oxidative lipid stress. In this context, HSD17B11 may promote a compartmentalized antioxidant system on LDs by maintaining FSP1 localization and stability, thereby protecting neutral lipids and limiting the propagation of lipid peroxidation.

In addition to regulating FSP1, HSD17B11 deficiency altered multiple LD features, including LD size and subcellular distribution. These phenotypes were not recapitulated by FSP1 deletion, suggesting that HSD17B11 functions beyond regulating FSP1 alone. Given the central role of LDs in lipid metabolism and oxidative stress responses^7,27,28^, our findings raise the possibility that HSD17B11 coordinates multiple aspects of LD biology, lipid metabolism, and antioxidant defense. Future studies investigating the molecular and metabolic functions of HSD17B11 may therefore provide important insights into how cells integrate lipid homeostasis with protection against oxidative stress.

HSD17B11 is closely related to the LD protein HSD17B13, a liver-enriched factor genetically associated with protection from metabolic liver disease^20^. However, unlike HSD17B13, HSD17B11 is broadly expressed across tissues^19^, suggesting that the HSD17B11-FSP1 pathway may represent a more fundamental and widely utilized mechanism for regulating compartmentalized ferroptosis defense. More broadly, our findings raise the possibility that LD-associated proteins may actively organize antioxidant machineries on LD surfaces to coordinate lipid metabolic homeostasis with protection against oxidative stress.

## MATERIALS AND METHODS

### Cell lines

The human hepatocellular carcinoma cell line Huh7 and human embryonic kidney-derived HEK293T cells were maintained in Dulbecco’s Modified Eagle Medium (DMEM) supplemented with 10% heat-inactivated fetal bovine serum (FBS), 25 mM HEPES, and MycoZap Plus-CL (Lonza) at 37°C in a humidified incubator with 5% CO_2_.

### Plasmid and siRNA transfection

Cells were plated 24 h before transfection. Plasmid transfection was performed using TransIT-LT1 reagent (Mirus Bio) according to the manufacturer’s instructions, with DNA amounts optimized for individual constructs. Cells were typically analyzed 24 h after transfection.

For siRNA-mediated knockdown, ON-TARGETplus SMARTpool siRNAs (Horizon Discovery) were delivered using TransIT-X2 reagent (Mirus Bio). Two rounds of transfection were performed at Day 1 and Day 3 after cell seeding, and cells were analyzed 6–7 days after plating.

### CRISPR/Cas9-mediated gene knockout

Knockout cell lines were generated using the CRISPR/Cas9 system. Guide RNA (gRNA) plasmids targeting HSD17B11 and FSP1 were obtained from Addgene. For HSD17B11 knockout, cells were transfected with CRISPR plasmid (Addgene #161923^29^) and seeded into 96-well plates for single-cell isolation. Candidate clones were screened by immunofluorescence to confirm loss of HSD17B11 expression.

For FSP1 knockout, lentiviral particles were produced in HEK293T cells by transfection with the corresponding gRNA plasmid (Addgene #186026^30^). Viral supernatants were collected 72 h post-transfection and used to infect Huh7 cells. Single-cell colonies were generated and screened by immunoblotting to confirm FSP1 deletion.

### Immunofluorescence and confocal microscopy

Cells cultured on coverslips were fixed with 4% paraformaldehyde (PFA) and washed three times with PBS. Primary antibodies were diluted in PBS containing 1% BSA and 0.2% saponin and incubated either overnight at 4°C or for 1 h at 37°C. Alexa Fluor-conjugated secondary antibodies (Thermo Fisher Scientific) were applied in the same buffer for 30 min at 37°C.

For lipid droplet staining, BODIPY 493/503 (10 μM; ThermoFisher) was added together with secondary antibodies. Coverslips were mounted using DAPI-containing mounting medium (Electron Microscopy Sciences) and dried at 37°C before imaging. Images were acquired using a Zeiss LSM900 confocal microscope equipped with ZEN software.

### High-content imaging and image analysis

Cells cultured in optical 96-well plates were fixed and stained as described above and maintained in PBS before imaging. Imaging and quantitative analysis were performed using the CellInsight high-content imaging platform (Thermo Fisher Scientific). DAPI staining was used for autofocus and nuclear segmentation, while CellMask staining defined cell boundaries. Cells exhibiting incomplete boundaries, abnormal morphology, or signs of cell death were excluded from analysis.

For LD quantification, cell-enclosed regions were defined as ROI_A. BODIPY-positive puncta were detected using the spot detection function, and puncta overlapping ROI_A were quantified based on fluorescence intensity or area. Colocalization analysis was performed using the overlap area output generated by the software colocalization function. To quantify peripheral LD distribution, an image analysis approach adapted from our previous lysosome positioning study^31^ was applied. Briefly, a peripheral ring region (ROI_B) was generated by shrinking the cell boundary inward by 15 pixels. The percentage of LDs localized within ROI_B relative to total cellular LDs was calculated as peripheral LD percentage.

### LD purification

LD purification was adapted from a previously published protocol. Briefly, approximately 3 × 10^8^ cells were harvested in PBS and disrupted in buffer A (20 mM tricine, 250 mM sucrose, pH 7.8, supplemented with protease inhibitors) using a nitrogen bomb at 35 bar after 15 min equilibration. Following centrifugation at 3,000 × g for 10 min to remove nuclei and unbroken cells, supernatants were subjected to ultracentrifugation with buffer B (20 mM HEPES, 100 mM KCl, 2 mM MgCl_2_, pH 7.4) layered above the lysate. After centrifugation at 182,000 × g for 1 h in a swinging-bucket rotor, LD fractions were collected and washed three times with buffer B. LD proteins were extracted in TBS containing 1% SDS, and residual lipids were removed by brief centrifugation.

### Cell fractionation

Approximately 2 × 10^7^ cells were harvested and mechanically disrupted by repeated passage through a 26-gauge needle in buffer A. LD fractions were isolated using the same flotation procedure described for LD purification. After collection of the LD fraction, the remaining lysate was subjected to ultracentrifugation at 270,000 × g for 1 h to separate cytosolic and membrane fractions. The cytosolic fraction was carefully collected from the middle layer of the tube using a loading tip. Membrane pellets were washed once with buffer B and solubilized in 2× SDS sample buffer by sonication.

### Immunoblotting

Cells were lysed directly in 2× SDS sample buffer, and lysates were subjected to heat-vortex cycles to break DNA. Protein samples were mixes with 4X SDS sample buffer. Samples were separated by SDS-PAGE and transferred to NC membranes followed by incubation with the indicated primary and HRP-conjugated secondary antibodies.

### Statistical analysis

Statistical significance was assessed using GraphPad Prism 9 (GraphPad Software). For comparisons between two groups, two-tailed t-*t*ests (paired or unpaired, as appropriate based on experimental design) were used. For comparisons involving more than two groups, one-way analysis of variance (ANOVA) was performed. All statistical tests were applied assuming approximately normally distributed data and similar variances between groups, as is standard for parametric analyses. The statistical significance is generally denoted as follows: *, *p* < 0.05, **, *p* < 0.01, ***, *p* < 0.001, ****, *p* < 0.0001, and n.s., not significant.

## ACKNOWLEDGMENTS

This research made use of the CellInsight Microscope Platform and confocal microscope from Autophagy, Inflammation, and Metabolism Center of Biomedical Research Excellence, funded by NIGMS, NIH (P20GM121176). Mass spectrometry analysis was performed at Taplin Biological Mass Spectrometry Facility, Harvard Medical School. The funding support was from NIGMS, NIH (T32GM144834 and R35GM147419).

## AUTHOR CONTRIBUTIONS

J.P. conceived the project. V.M.V. performed most of the experiments. B.G. performed cell death assays. H.A. performed immunoblotting for cell fractionation assays. O.K. generated the HSD17B11-KO cells. L.C. measured lipid droplet distribution. N.N. and J.R. quantified lipid droplet size distribution. M.R.L. performed AlphaFold analysis. V.M.V. and J.P. analyzed the data. J.P. wrote the manuscript. All the authors edited the manuscript.

## DECLARATION OF GENERATIVE AI AND AI-ASSISTED TECHNOLOGIES IN THE MANUSCRIPT PREPARATION PROCESS

During the preparation of this work the authors used ChatGPT in order to assist with grammar checking and language refinement. After using this tool/service, the authors reviewed and edited the content as needed and takes full responsibility for the content of the published article.

## REFERENCES

1. Dixon, S. J. et al., Stockwell, B. R. Ferroptosis: an iron-dependent form of nonapoptotic cell death. Cell 149, 1060–1072 (2012).

2. Lei, G., Zhuang, L. & Gan, B. Targeting ferroptosis as a vulnerability in cancer. Nat Rev Cancer 22, 381–396 (2022).

3. Wahida, A. & Conrad, M. Decoding ferroptosis for cancer therapy. Nat Rev Cancer 25, 910–924 (2025).

4. Belavgeni, A., Meyer, C., Stumpf, J., Hugo, C. & Linkermann, A. Ferroptosis and Necroptosis in the Kidney. Cell Chem Biol 27, 448–462 (2020).

5. Ryan, S. K. et al., Hammond, T. R. Therapeutic inhibition of ferroptosis in neurodegenerative disease. Trends Pharmacol Sci 44, 674–688 (2023).

6. Jiang, X., Stockwell, B. R. & Conrad, M. Ferroptosis: mechanisms, biology and role in disease. Nat Rev Mol Cell Biol 22, 266–282 (2021).

7. Dixon, S. J. & Olzmann, J. A. The cell biology of ferroptosis. Nature reviews Molecular cell biology 25, 424–442 (2024).

8. Yang, W. S. et al., Stockwell, B. R. Regulation of ferroptotic cancer cell death by GPX4. Cell 156, 317–331 (2014).

9. Doll, S. et al., Conrad, M. FSP1 is a glutathione-independent ferroptosis suppressor. Nature 575, 693–698 (2019).

10. Bersuker, K. et al., Olzmann, J. A. The CoQ oxidoreductase FSP1 acts parallel to GPX4 to inhibit ferroptosis. Nature 575, 688–692 (2019).

11. Dierge, E. et al., Feron, O. Peroxidation of n-3 and n-6 polyunsaturated fatty acids in the acidic tumor environment leads to ferroptosis-mediated anticancer effects. Cell Metab 33, 1701–1715.e5 (2021).

12. Lee, H. et al., Gan, B. Cell cycle arrest induces lipid droplet formation and confers ferroptosis resistance. Nat Commun 15, 79 (2024).

13. Herms, A. et al., Pol, A. AMPK activation promotes lipid droplet dispersion on detyrosinated microtubules to increase mitochondrial fatty acid oxidation. Nat Commun 6, 7176 (2015).

14. Motamedi, S. et al., Swinnen, J. V. AMP-activated protein kinase-driven lipid droplet dynamics govern melanoma sensitivity to polyunsaturated fatty acid and iron-induced ferroptosis. Nat Commun 16, 11252 (2025).

15. Chang, C.-L. et al., Lippincott-Schwartz, J. Spastin tethers lipid droplets to peroxisomes and directs fatty acid trafficking through ESCRT-III. J Cell Biol 218, 2583–2599 (2019).

16. Kong, J. et al., Kim, J. B. Spatiotemporal contact between peroxisomes and lipid droplets regulates fasting-induced lipolysis via PEX5. Nat Commun 11, 578 (2020).

17. Lange, M. et al., Olzmann, J. A. FSP1-mediated lipid droplet quality control prevents neutral lipid peroxidation and ferroptosis. Nat Cell Biol 27, 1902–1913 (2025).

18. Nakamura, T. et al., Conrad, M. Phase separation of FSP1 promotes ferroptosis. Nature 619, 371–377 (2023).

19. Liang, B., Fu, L. & Liu, P. Regulation of lipid droplet dynamics and lipid homeostasis by hydroxysteroid dehydrogenase proteins: (Trends in Cell Biology, published online November 26, 2024). Trends Cell Biol 36, 175–176 (2026).

20. Abul-Husn, N. S. et al., Dewey, F. E. A Protein-Truncating HSD17B13 Variant and Protection from Chronic Liver Disease. N Engl J Med 378, 1096–1106 (2018).

21. Deol, K. K. et al., Olzmann, J. A. Vitamin B2 metabolism promotes FSP1 stability to prevent ferroptosis. Nat Struct Mol Biol 33, 525–536 (2026).

22. Horiguchi, Y., Araki, M. & Motojima, K. Identification and characterization of the ER/lipid droplet-targeting sequence in 17beta-hydroxysteroid dehydrogenase type 11. Arch Biochem Biophys 479, 121–130 (2008).

23. Wu, K. et al., Papagiannakopoulos, T. Targeting FSP1 triggers ferroptosis in lung cancer. Nature 649, 487–495 (2026).

24. Yoshioka, H. et al., Osada, H. Identification of a Small Molecule That Enhances Ferroptosis via Inhibition of Ferroptosis Suppressor Protein 1 (FSP1). ACS Chem Biol 17, 483–491 (2022).

25. Hendricks, J. M. et al., Olzmann, J. A. Identification of structurally diverse FSP1 inhibitors that sensitize cancer cells to ferroptosis. Cell Chem Biol 30, 1090–1103.e7 (2023).

26. Lee, H. et al., Gan, B. Cell cycle arrest induces lipid droplet formation and confers ferroptosis resistance. Nat Commun 15, 79 (2024).

27. Farese, R. V. & Walther, T. C. Essential Biology of Lipid Droplets. Annu Rev Biochem 94, 447–477 (2025).

28. Henne, W. M. & Cohen, S. Heterogeneity, dynamics and organelle interactions of lipid droplets. Nat Rev Mol Cell Biol (2026).

29. Demange, P. et al., Britton, S. SDR enzymes oxidize specific lipidic alkynylcarbinols into cytotoxic protein-reactive species. Elife 11, e73913 (2022).

30. Mao, C. et al., Gan, B. DHODH-mediated ferroptosis defence is a targetable vulnerability in cancer. Nature 593, 586–590 (2021).

31. Kolaczkowski, O. M. et al., Pu, J. Synergistic Role of Amino Acids in Enhancing mTOR Activation Through Lysosome Positioning. bioRxiv 2024.10.12.618047 (2024).

